# KHDRBS3 facilitates self-renewal and temozolomide resistance of glioblastoma cell lines

**DOI:** 10.1101/2024.06.16.599185

**Authors:** Kanokkuan Somrit, Sucheewin Krobthong, Yodying Yingchutrakul, Nut Phueakphud, Patompon Wongtrakoongate, Waraporn Komyod

## Abstract

Glioblastoma is a deadly tumor which possesses glioblastoma stem cell populations involved in temozolomide resistance. To gain insight into the mechanisms of self-renewing and therapy-resistant cancer stem cells, subcellular proteomics was utilized to identify proteins whose expression is enriched in U251-derived glioblastoma stem-like cells. The RNA binding protein KHDRBS3 was successfully identified as a gene up-regulated in the cancer stem cell population compared with its differentiated derivatives. Depletion of KHDRBS3 by RNA silencing led to a decrease in cell proliferation, neurosphere formation, migration, and expression of genes involved in glioblastoma stemness. Importantly, temozolomide sensitivity can be induced by the gene knockdown. Collectively, our results highlight KHDRBS3 as a novel factor associated with self-renewal of glioblastoma stem-like cells and temozolomide resistance. As a consequence, targeting KHDRBS3 may help eradicate glioblastoma stem-like cells.

## Introduction

Glioblastoma is the most aggressive primary brain cancer. Temozolomide is a first-line cytotoxic drug treatment for glioblastoma that prolongs overall survival with a median survival of approximately 15 months (McNamara et al., 2014, Norden et al., 2010). Despite recent therapeutic advances, including the combination of surgical resection, radiotherapy and chemotherapy, less than one-third of glioblastoma patients survive more than two years. Moreover, patients with glioblastoma show highly variable responses to chemotherapy, and tumor recurrence is frequently observed (Jiapaer et al., 2018, Wu et al., 2021). Resistance and subsequent tumor relapse of glioblastoma patients could be driven by the presence of therapy-resistant, genetically distinct tumor subpopulations. Novel targeted therapies for glioblastoma are needed to achieve a complete remission of the cancer.

Glioblastoma stem cells are subpopulations of glioblastoma, and have been proposed to be responsible for tumor initiation, therapeutic resistance and recurrence (Auffinger et al., 2015, Biserova et al., 2021, Bradshaw et al., 2016, Lah et al., 2020). Furthermore, cancer stem cells have been suggested to be the source of the metastatic trait. In colorectal cancer, metastasis can be induced by injection of a tumor population containing colon cancer stem cells into an animal model (Pang et al., 2010). Regulation of glioblastoma stem cells involves both extrinsic and intrinsic factors. Extrinsically, for example, glioblastoma stem cells maintain stem cell state by aberrant activation of common signaling pathways such as Notch, Wnt, and PI3K-AKT signaling. Intrinsically, to resist to temozolomide, the stem cell population in glioblastoma can carry promoter methylation of the gene *MGMT* encoding the DNA repair enzyme O^6^-methylguanine-DNA methyltransferase resulting in an induction of MGMT expression, which protects the genome from the cytotoxic action of the chemotherapeutic drug (Biserova et al., 2021, Lathia et al., 2015, Tomar et al., 2021). Unfortunately, almost all patients with glioblastoma relapse after surgery and chemotherapy due to residual tumors which can be enriched for the cancer stem cells leading to even greater resistant against temozolomide (Chen et al., 2012, Putavet and de Keizer, 2021). Therefore, an identification of novel factors facilitating stem cell maintenance of glioblastoma and temozolomide resistance is crucial to design next-generation anti-cancer treatments.

In this study, we aim to identify novel regulators involved in glioblastoma stem cells and temozolomide resistance. Through nuclear proteomic analyses of glioblastoma stem-like cells and their differentiated counterparts, we identify the protein KHDRBS3 as a key mediator of cancer stem cell maintenance and resistance to temozolomide of glioblastoma cell lines. The results presented here reveal the role of KHDRBS3 in regulation of cancer phenotypes associated with glioblastoma stem-like cells including temozolomide resistance, which will help to improve the molecular understanding of glioblastoma stem cells and the development of anti-glioblastoma therapy.

## Methods

### Cell culture and small interfering RNA (siRNA) transfection

Human GBM cell lines U251, LN229, U87 and M059J were maintained in Dulbecco’s modified Eagle’s medium/Ham’s F-12 medium (DMEM/F-12) (HyClone, SH30023.02) supplemented with 10% fetal bovine serum (FBS) (Thermo Fisher Scientific, 10270-106) at 37 ^◦^C in a 5% CO_2_ humidified incubator and passaged every 3-4 days. KHDRBS3 knockdown was performed by transfection of 50 nM of non-targeting siRNA (si-CTL) or specific siRNA targeting KHDRBS3 (si-KHDRBS3: 5′ GCGAAGUACUUCCAUCUCAAUGA-3’) (Dharmacon) with Lipofectamine RNAiMAX Transfection Reagent (Invitrogen, #13778075) following manufacturer’s instructions. U251 cells were seeded at density 5 x 10^5^ cells/mL 24 h before siRNA transfection. After 24 h post-transfection, cells were subsequently analyzed or further cultured under spheroid condition.

### Neurosphere formation assay

Cells were trypsinized with 0.25% Trypsin-EDTA (Thermo Fisher Scientific, 25200056) and seeded at the density of 5 x 10^3^ cells/mL in non-treated 48 well-plate in DMEM/F-12 supplemented with 20 ng/mL epidermal growth factor (EGF) (Peprotech, AF-100-15), 20 ng/mL basic fibroblast growth factor (bFGF) (Prepotech, AF-100-18B) and 1x B27 (Thermo Fisher Scientific, 17504044). Cells were incubated at 37°C in a humidified 5% CO_2_ atmosphere, and the fresh culture medium was added every three days without removing remaining media. The neurospheres with a diameter > 100 μm were counted under microscopy and expressed as mean ± SEM. After 7 days, neurospheres were collected or attached to standard culture plates in media containing 5% FBS for 6 h and stained with crystal violet solution (Sigma-Aldrich, 548-62-9).

### Sodium butyrate treatment

Sodium butyrate (Sigma-Aldrich, QU6005-01265) was dissolved in H_2_O. To induce differentiation, neurospheres at day 7 were collected and dissociated into single cells by trypsinization. Cells were seeded at 10^4^ cells/cm^2^ in a normal culture dish in culture media containing 3 mM of SB for 7 days. Culture media containing SB was changed every 2 days.

### Subcellular cytoplasmic and nuclear protein extraction

Nuclear and cytoplasmic proteins were extracted from the parental, differentiated and spheroid cells at day 7. Briefly, cells were washed twice with PBS and incubated in cytoplasmic extract buffer (10 mM Hepes-KOH, pH 7.9, 1.5 mM MgCl_2_, 10 mM KCl) supplemented with freshly 1 mM Na_3_VO_4_, 1 mM PMSF, 1mM DTT and protease inhibitor, and incubated on ice for 15 min. After adding 10% Nonidet P-40, vortexed for 5 seconds, cells were centrifuged at 4°C 2000 rpm for 1 min. Supernatant was carefully transferred to a centrifuge tube and collected as cytoplasmic protein. The pellet was resuspended in 50 µl high-salt buffer (20 mM Hepes-KOH, pH 7.9, 1.5 mM MgCl_2_, 10 mM KCl, 420 mM NaCl, 0.2 mM EDTA and 25% Glycerol) supplemented with freshly 1mM Na_3_VO_4_, 1 mM PMSF, 1mM DTT and protease inhibitor, and incubated on ice for 30 min. After centrifugation at 13000 rpm for 10 min, the supernatant was collected as nuclear protein and stored at −80 °C. The concentration and purity of extracted protein was determined using Bio-Rad Protein Assay Dye Reagent Concentrate (Bio-Rad, 500-0006) following manufacturer’ s protocol.

### Sample preparation and protein digestion

Equal amounts of proteins (50 μg) from parental, differentiated and spheroid were reduced before trypsinization. Briefly, 200 mM DTT was added, and samples were incubated for 45 min to reduce the sample. The cysteine groups were blocked with 1 M iodoacetamide in the dark for 45 min. Trypsin (mass spectrometry grade; Promega, V5280) was used at a ratio of 1:30 (sample:enzyme) and incubated overnight at 37 °C. The reaction was stopped with the addition of 1% formic acid (Merck, 64-18-6) and peptides were dried in a SpeedVac, diluted in 0.1% formic acid, dried again and kept at −20 °C before LC-MS/MS analysis.

### Nano-liquid chromatography with mass spectroscopy

Peptides were fractionated by an ultimate 3000 nanoLC-MS/MS system (Thermo Scientific^TM^) using an analytical column C18 (reverse phase chromatography) (Thermo Fisher Scientific, 17126-032130) at 300nL/minute over a 145-minute gradient. Data-dependent LC-MS-MS was performed using High resolution 6600+ TripleTOF^TM^ (ABSciex). All samples were subjected to 1 DIA LC-MS run for generation of spectral ion library followed by 4 instrument replications (n=4) LC-MS runs for DDA (SWATH-MS) analysis. The MS was acquired using IDA and SWATH^TM^ acquisition data collection. Data dependent acquisition was performed in positive ion mode. The 50 most abundant ions of which charge states were from 2+ to 5+ were selected for subsequent fragmentation with rolling collision energy. MS/MS spectra were acquired from m/z 300 to m/z 1800.

### Protein identification and database search

The peptide data obtained from 6600+ TripleTOF^TM^ was analyzed with ProteinPilot^TM^ 3.0 software using the Paragon protein database search algorithm (AB Sciex). IDA data files from ProteinPilot^TM^ and SWAT MS from SWAT MicroApp 2.0 were subjected to isotope pattern matched peak mining using the Extracted Ion Chromatogram (XIC) manager add-on for PeakView. Peptide MS and retention time were calibrated using MarkerView software (AB Sciex). Database interrogation was: taxonomy – homo sapiens; database – Paragon: fixed modification – carbamidomethyl; variable modification - NQ deamidation, M oxidation, PK hydroxylation; precursor charge state – 2+ to 5+. The false discovery rate was below 1% for protein identification. For protein quantitation, ‘NormalyzerDE software’ was used for normalization. The list of differentially expressed proteins, including fold changes, was imported as an Excel file. Proteins with a fold change of two were generally considered as differentially expressed. Significant differences in protein expression levels were determined by Student’s t-test with a set value of P<0.05. The gene ontology (GO) and pathway analyses were performed with open-source software ShinyGO (version 0.76.2, http://bioinformatics.sdstate.edu/go/) selected Homo sapiens organisms. For determination of protein interaction networks, the list of differentially expressed proteins was imported in the online STRING database (https://string-db.org) of known and predicted protein interactions. The interactions include direct (physical) and indirect (functional) associations between proteins.

### Quantitative realtime PCR

Total RNA was extracted by illustra^TM^ RNAspin Mini Isolation RNA kit (Cytiva, GE25-0500-71) according to the manufacturer’s instructions. The first strand cDNA was synthesized from 1 ug of total RNA using HyperScript^TM^ reverse transcription system (GeneAll, 601-710). Quantitative real-time PCR was performed on Bio-Rad CFX96 Real-Time PCR Detection System with SYBR Green Master Mix (PCR Biosystems, PB20.11-05). Three independent quantitative RT-PCR experiments were done in triplicate. Results were normalized to the threshold cycle (Ct) of internal control *β-actin*, referred to as △^Ct^. The fold change was presented as 2^△△Ct^. PCR primer sequences are available upon requested.

### Flow cytometry

Cells were collected with 0.05% Trypsin-EDTA to preserve the surface antigens and resuspended in Flow staining buffer (1% w/v BSA and 0.25 mM EDTA pH 8 in PBS) 100 μL per 10 cells. To determine the cancer stem-like subpopulations in spheroids, differentiated and parental, the expression of the surface marker CD133 was determined using CD133 conjugated allophycocyanin (APC) antibody (Miltenyi, 130-113-668, 1:50 dilution) for 10 minutes in the dark at 4 °C followed by washing twice before acquisition. Annexin V-FITC (Biovision, 1001) and propidium iodine (PI) (Invitrogen, P3566) double staining were used to determine apoptotic cell death according to the manufacturer’s protocol. Then, flow cytometry was performed with a FACSCanto II Flow Cytometer (BD Biosciences). Data were analyzed and presented using BD FACSCanto II System Software (version 3.0).

### Drug sensitivity assay

The sensitivity of parental differentiated and spheroids to temozolomide (TEMOZOLOMIDE) was measured by MTT-based cell viability assay. Spheroid differentiated and parental cells were seeded at a density of 5000 cells/well in 96-well plates and cultured in normal culture medium. Different concentrations of TEMOZOLOMIDE (Sigma-Aldrich, T2577) ranging from 0 μM to 400 μM and DMSO alone (the controls) were added to the cell cultures in a total volume of 200 μL/well. After 48 or 72 hours of incubation, the assay was performed by adding 100 μL of 3-(4, 5-dimethyl-2-thiazolyl)-2, 5-diphenyl-2H-tetrazolium bromide solution (MTT) (Sigma-Aldrich, A2231, 5 mg/mL in PBS) to each well, incubating 3 hours, adding 200 μL of DMSO to dissolve the formazan crystals, and reading the absorbance of each well at 540 nm by a microplate reader. The half-maximal inhibitory concentration (IC50) was determined using the Chou-Talalay method. The cytotoxicity was determined by comparing the resulting absorbance with the mean absorbance of the control wells (considered as 100% viability) and was expressed as percentage of cell viability.

### Western blot analysis

Lysates of spheres and parental were prepared in lysis buffer (20 mM HEPES pH7.4, 1% Nonidet P-40, 150 mM NaCl, 5 mM MgCl_2_, 10% glycerol) with protease inhibitor (Roche Diagnostics, 4693132001). The protein samples (40 μg) were separated by electrophoresis on 10% polyacrylamide SDS gels (120 V for 1 to 2 h) and transferred to polyvinylidene difluoride (PVDF) membrane (PALL, BSP0861) using Semi-dry electroblotting (0.5 A for 1 h). Then, non-specific binding was blocked by incubating the membrane with 5% bovine serum albumin (BSA) in TBS-T (20 mM Tris, 137 mM NaCl, 0.1% Tween 20) for 1 h. The membrane was rinse twice with TBS-T and probed with KHDRBS3 rabbit polyclonal antibody (Proteintech, 13563-1-AP, 1:1000 dilution) or β-actin mouse polyclonal antibody (Sigma-Aldrich, A1978,1:1000 dilution) at 4°C overnight. After incubation with horseradish peroxidase (HRP)-conjugated anti-rabbit or anti-mouse secondary antibodies (Abcam, ab205719, ab205718, 1:1000 dilution) for 45 min at room temperature, the blot was detected using Clarity^TM^ ECL Western blotting substrate (Bio-Rad, 1705061). The intensity of bands will be quantified using Image J software (NIH image, National Institutes of Health, Bethesda, MD, USA). All Western blot analyses were repeated at least three times.

### Wound healing assay

For migration assays, si-CTL and si-KHDRBS3 treated cells were seeded at density of 2.5 x 10^5^ cells/well and cultured to confluence for 24 h. The cell monolayer was then scraped with a pipette tip (t = 0 h) and cultures were monitored at 24 h to evaluate their wound healing ability. Images were captured and migration rates were measured using ImageJ software (NIH, Bethesda, MD) and expressed as percentage of migration compared to that at 0 h.

### Statistical analysis

All experiments were repeated independently at least three times. Data were analyzed and represented as mean values ± standard deviation (SD). Statistical comparisons were made with a two-tailed unpaired Student’s t-test. A P-value of less than 0.05 was considered as significant.

## Results

### Glioblastoma stem-like cells can be enriched in U251 neuropheres

Among four glioblastoma cell lines, we observed a highest expression of key neural stem-associated genes in U251, suggesting its potential to represent a model of glioblastoma stem cells (Figure S1). To ascertain whether the human glioblastoma cell line U251 can be utilized as the model of glioblastoma stem cells, the cell line was shifted from a serum-containing condition routinely used to maintain the cells, to a serum-free condition containing B27, EGF and bFGF to facilitate self-renewal (Qiang et al., 2009b). Under this neurosphere-forming condition, U251 began to form spheroids on day 2 (Figure 1A). The size of the spheroids could also be expanded on days 4 and 6. Realtime PCR analysis confirmed that expression of the neural stem cell gene *SOX2* is induced in the spheroids (Figure 1C). On the other hand, expression of the glial cell differentiation maker *GLAST* is decreased in the spheroids. As determined by flow cytometry, these neurosphere cells up-regulate expression of the cell surface antigen CD133, which has been used as a marker of glioblastoma cancer stem cells (Figure 1D) (Brown et al., 2017, Jin et al., 2011). Further, to induce differentiation of U251, cells from the spheroids were dissociated, and subsequently treated with the differentiation-inducing agent sodium butyrate (SB) for 6 days. As expected, the cells elongate and expand in size upon treatment with sodium butyrate (Figure 1B). In addition, while expression of the neural stem cell genes SOX2 and PAX6 was reduced, that of GLAST was induced in the sodium butyrate-treated cells (Figure 1C). Moreover, the number of cells positive for CD133 was reduced compared with cells grown under the serum-containing condition and cells grown under the spheroid condition (Figure 1C). These results suggest that the stem cell-like state of the glioblastoma cells U251 can be sustained by the culture condition used for maintenance of neural stem cells.

**Figure 1.**
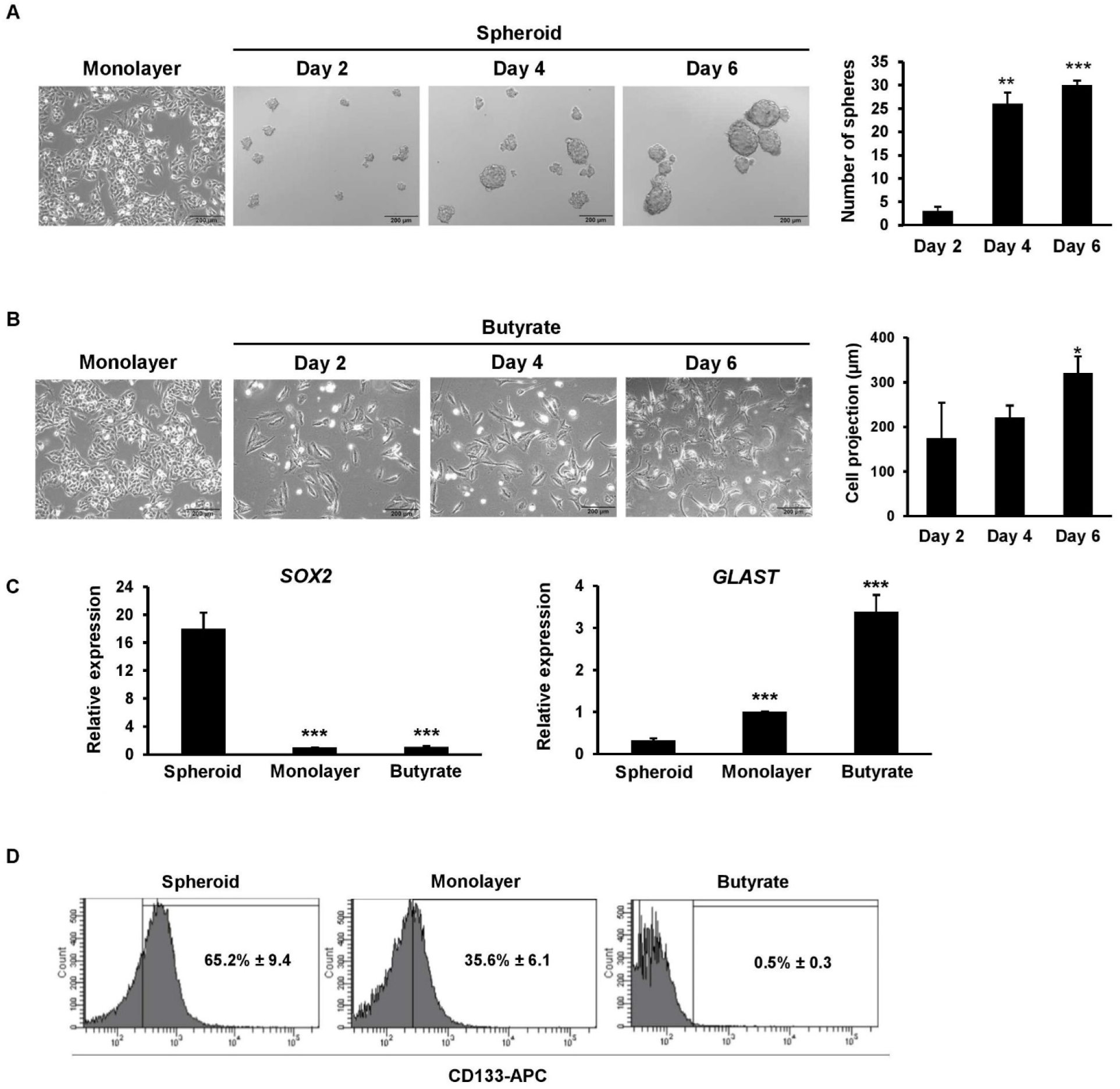
Glioblastoma stem-like and differentiated phenotypes can be induced by culture conditions. (A) U251 cells were seeded at a density of 5 x 10^3^ cells/mL in non-treated 48 well-plate to induce neurosphere formation. The morphology of U251 monolayer on day 6 and spheroid culture on day 2, 4 and 6 were observed under bright field (magnification: 10X, scale bar: 200 μm). Sphere size was observed every 2 days until day 6. Representative images of spheres are shown on the left, and sphere-forming efficiency is shown on the right. (B) Representative morphology of monolayer on day 6 and sodium butyrate-induced differentiated U251 on day 2, 4 and 6 are shown on the left. The average of cell projection is shown on the right. * p < 0.05, ** p < 0.01 and *** p < 0.005. (C) Realtime PCR analysis of stemness- and differentiation-related related genes *SOX2* and *GLAST* was performed using RNA collected from spheroid, monolayer, and butyrate-treated U251 cells. *ACTB* was used as internal control. The relative value was expressed as the mean ± SD from three independent experiments. *** p < 0.005. (D) The percentage of CD133-positive cells was determined using flow cytometry. The number represents the percentage of CD133-positive cells from three independent experiments.

### U251 neurospheres are resistant to the drug temozolomide

Cancer stem cells can often be characterized by their ability to desensitize chemotherapeutic agents (Phi et al., 2018, Wongtrakoongate, 2015). To determine whether the spheroids derived from U251 possess an increase in chemoresistance ability toward the conventional glioblastoma medication temozolomide, the sensitivity of cells grown under the spheroid, serum-containing and sodium butyrate-inducing conditions was compared for their half-maximal inhibitory concentration (IC50). Cells in each condition were cultured under serially diluted temozolomide. We observed a significant 2.2-fold (360±24 versus 163±15) increase in IC50 value comparing between the spheroid and the serum-containing conditions. In addition, a significant 3.5-fold (360±24 versus 103±14) increase in IC50 value was obtained by comparing between the spheroid and the sodium butyrate-containing conditions (Figure 2). This result indicates that the glioblastoma cell line U251 enriched for the glioblastoma stem-like cells in the neurosphere condition exhibits a higher chemoresistance ability than the two other conditions possessing less glioblastoma stem-like cells.

**Figure 2.**
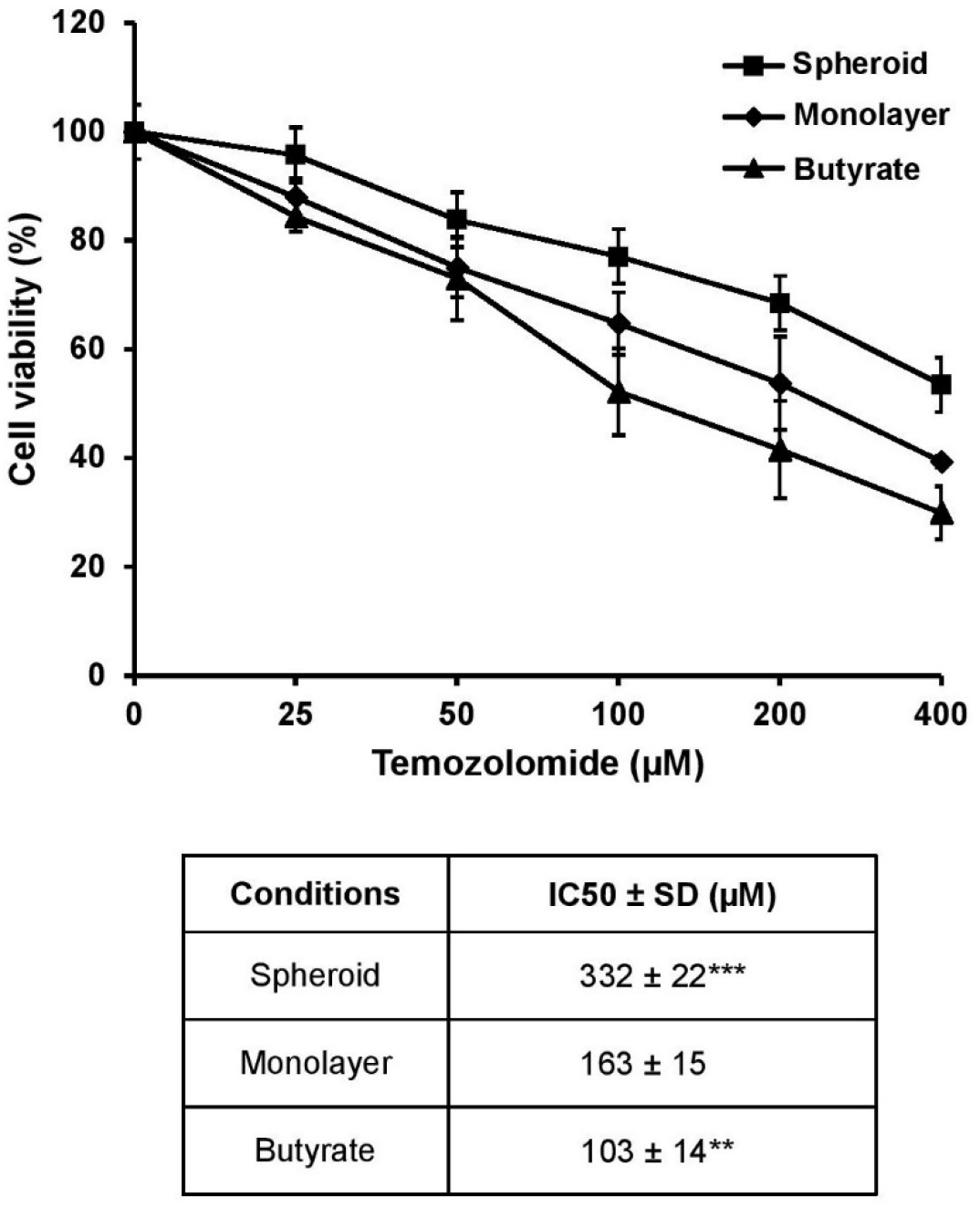
The neurosphere culture of U251 possesses the highest drug resistance against temozolomide. Cells derived from spheroid, monolayer, and butyrate-treated U251 cultures were seeded in 96-well plates and treated with the indicated concentrations of temozolomide for 72 h. Cell viability was determined by MTT assay. Error bars represent SD (n=3). The table shows the IC50 values (µM). All data are expressed as mean ± SD from three independent experiments. ** p < 0.01 and *** p < 0.005 vs monolayer culture.

### Up-regulation of the RNA-binding protein KHDRBS3 in U251 neurospheres

Next, we asked which proteins are differentially expressed under spheroid, monolayer, and butyrate-inducing conditions by performing nuclear and cytoplasmic proteomic analyses. Fractionated protein lysates collected from the three culture conditions were prepared and subjected to nanoflow liquid chromatography (nano-LC) coupled with high-resolution tandem mass spectrometry (MS). Of the cytoplasmic proteomes, we identified 1,098 common proteins being expressed across the three culture conditions. Pathway enrichment analysis of biological processes shows that functions of cytoplasmic proteins up-regulated in the spheroid condition are enriched in mesenchyme migration and cellular physiology/metabolisms (Figure S2).

Of the nuclear proteome, a total of 1,481 nuclear proteins were identified in the three different culture conditions. By applying the criteria of greater than two-fold change and p-value < 0.05, 257 and 215 nuclear proteins were identified as significantly up-regulated and down-regulated in U251 neurospheres when compared to butyrate-inducing condition. In our analysis, we categorized the nuclear proteins based on their biological functions using ShinyGo software. Analysis of biological processes using the up-regulated spheroid nuclear proteins indicated a strong involvement in the mRNA processing and RNA splicing, as well as DNA and RNA metabolomic processes. Up-regulated nuclear proteins enriched in the molecular functions of RNA binding, chromatin binding, and mRNA binding (Figures 3A-3B). To improve our understanding of the pathways involving the differentially expressed nuclear proteins, the KEGG analysis was performed. The KEGG pathway analysis of up-regulated nuclear proteins in spheroids revealed an important representation of the spliceosome pathway (Figure 3C). By querying the list of nuclear proteins up-regulated in spheroids compared with monolayer culture, five major clusters including regulation of replication and transcription, gene expression (transcription), pre-mRNA processing, transport of mature mRNA, and cell cycle were identified using the STRING database (Figure 3D).

**Figure 3.**
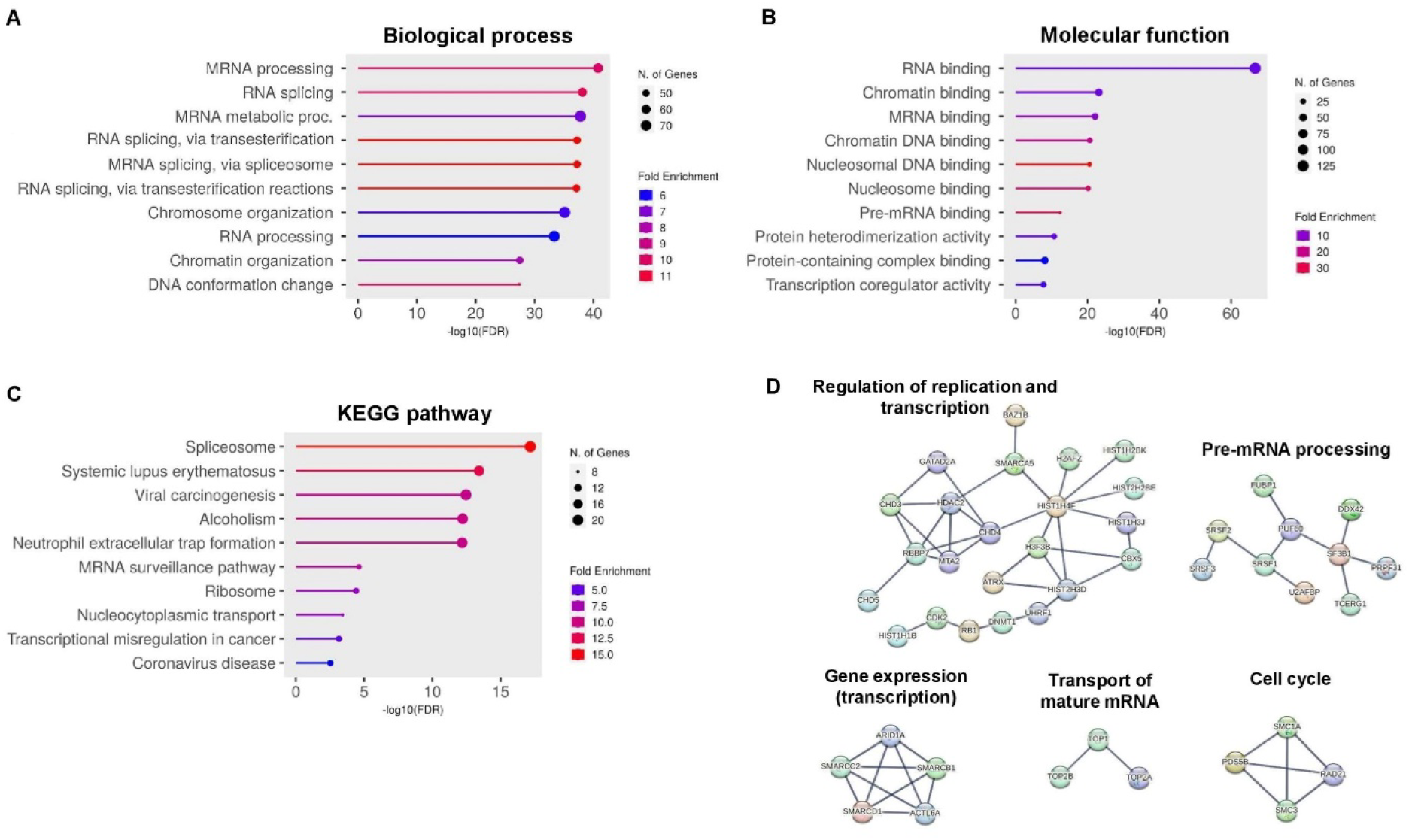
Differential nuclear proteomics of neurospheres and butyrate-treated U251 cells identifies proteins whose functions are associated with RNA biogenesis in glioblastoma stem-like cells. Top ten categories identified by gene ontology enrichment analysis of significant up-regulated proteins in spheroid vs butyrate-treated U251 according to (A) cellular processes, (B) molecular function and (C) KEGG pathway using ShinyGo software v0.76.2. The color chart shows the fold enrichment for each pathway. The size of dots corresponds to the number of genes assigned to each pathway. (D) Functional interactions of up-regulated nuclear proteins in the spheroids were identified using STRING. Networks with three or more protein interactions are shown. Required confidence (score) of protein association was 0.900 (highest confidence).

Expression of eight nuclear proteins (ARPC5, BAG3, HIX, HP1BP3, IF16, KHDRBS3, SPATS2L and TRIM22) and one cytoplasmic protein (CNTRL) was found to be at least two-fold up-regulated and down-regulated in neurospheres and butyrate-inducing condition compared with monolayer culture, respectively (Figure 4). Of the top two up-regulated proteins in neurospheres, BAG3 (BCL-2-intreacting cell death suppressor) is a well-characterized oncogene in glioblastoma. On the other hand, KH RNA binding domain containing, signal transduction associated 3 (KHDRBS3; also known as SLM2 or T-STAR) is an RNA-binding protein involved in RNA biogenesis, however the role of KHDRBS3 in glioblastoma biology and drug resistance has never been described. Realtime PCR was performed to ascertain whether expression of *KHDRBS3* and *BAG3* transcripts is up-regulated in neurospheres compared with monolayer cultures of the four glioblastoma cell lines including U251, LN229, U87, and M059J. The result shows that neurospheres derived from the four cell lines expressed KHDRBS3 and BAG3 higher than cells maintained in the serum-containing monolayer condition (Figures 4B-4C). Notably, the differential gene up-regulation is more prominent for U251 compared to other cell lines. Together, these results suggest that the glioblastoma neurospheres up-regulate KHDRBS3 and BAG3.

**Figure 4.**
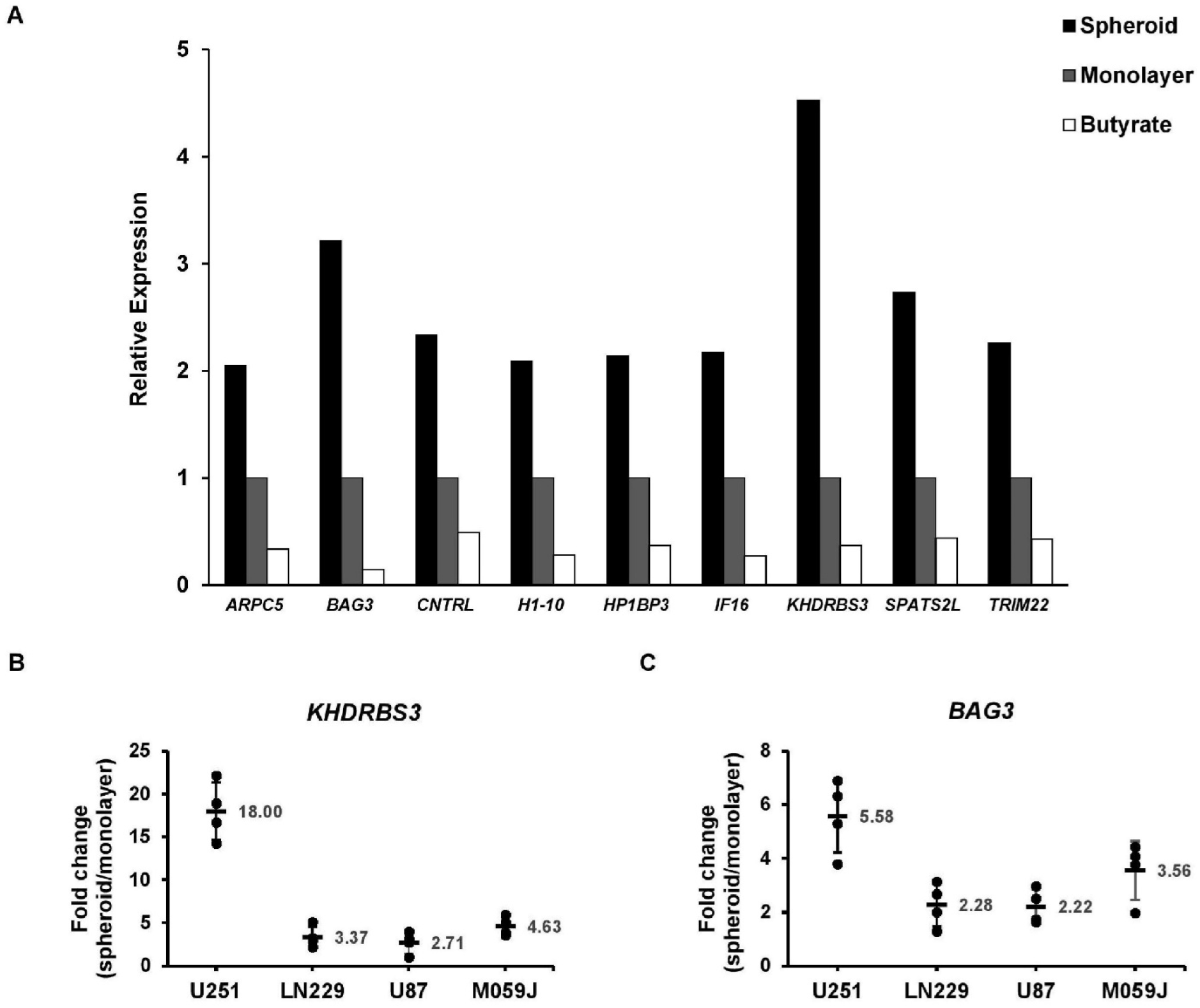
Expression of the gene encoding KHDRBS3 is up-regulated in glioblastoma-derived neurospheres. (A) Relative expression levels of nuclear proteins with at least two-fold changes and p-value < 0.05. The differential expression levels were shown for proteins up-regulated in spheroids over monolayer culture and for proteins down-regulated in butyrate treatment over monolayer culture. (B and C) Realtime PCR was performed to validate the up-regulation of *KHDRBS3* and *BAG3* in neurospheres of four glioblastoma cell lines including U251, LN229, U87 and M059J. The numbers indicate mean of fold-change. *ACTB* was used as internal control. The relative value was expressed as the mean ± SD (n=4). *** p < 0.005.

### KHDRBS3 facilitates self-renewal, migration, and temozolomide resistance

Given that expression of KHDRBS3 is highly induced in the neurospheres, we next elucidated the significance of KHDRBS3 up-regulation in self-renewal of U251. To this end, expression of KHDRBS3 was suppressed by using small interfering RNA (siRNA). U251 cells were transfected with control siRNA (si-CTL) or siRNA against KHDRBS3 (si-KHDRBS3) for 48 h, followed by incubation under the sphere-forming condition. As shown in Figures 5A and 5B, silencing of KHDRBS3 led to an efficient down-regulation of KHDRBS3 mRNA and protein expression. Similar results were also obtained from the cell line LN229. We then determined the effect of KHDRBS3 knockdown on cell proliferation. KHDRBS3 knockdown significantly decreased cell number and viability on day 6 (Figures 5C and 5D). As soon as day 3 upon sphere formation, KHDRBS3 depletion reduced the number of spheres with the size equal to or larger than 50 μ in both U251 and LN229 cell lines (Figure 6), suggesting that KHDRBS3 is important for the neurosphere formation in the glioblastoma stem-like cells. In addition, *in vitro* wound healing assay showed that KHDRBS3 suppression resulted in a 2-fold decrease in migration ability of U251 cells (Figure 7). Moreover, realtime PCR analysis revealed that silencing of *KHDRBS3* led to a down-regulation of genes involved in self-renewal of glioblastoma stem-like cells including *SOX2*, *PAX6*, *CD133*, and *CD44* transcripts (Figure 8).

**Figure 5.**
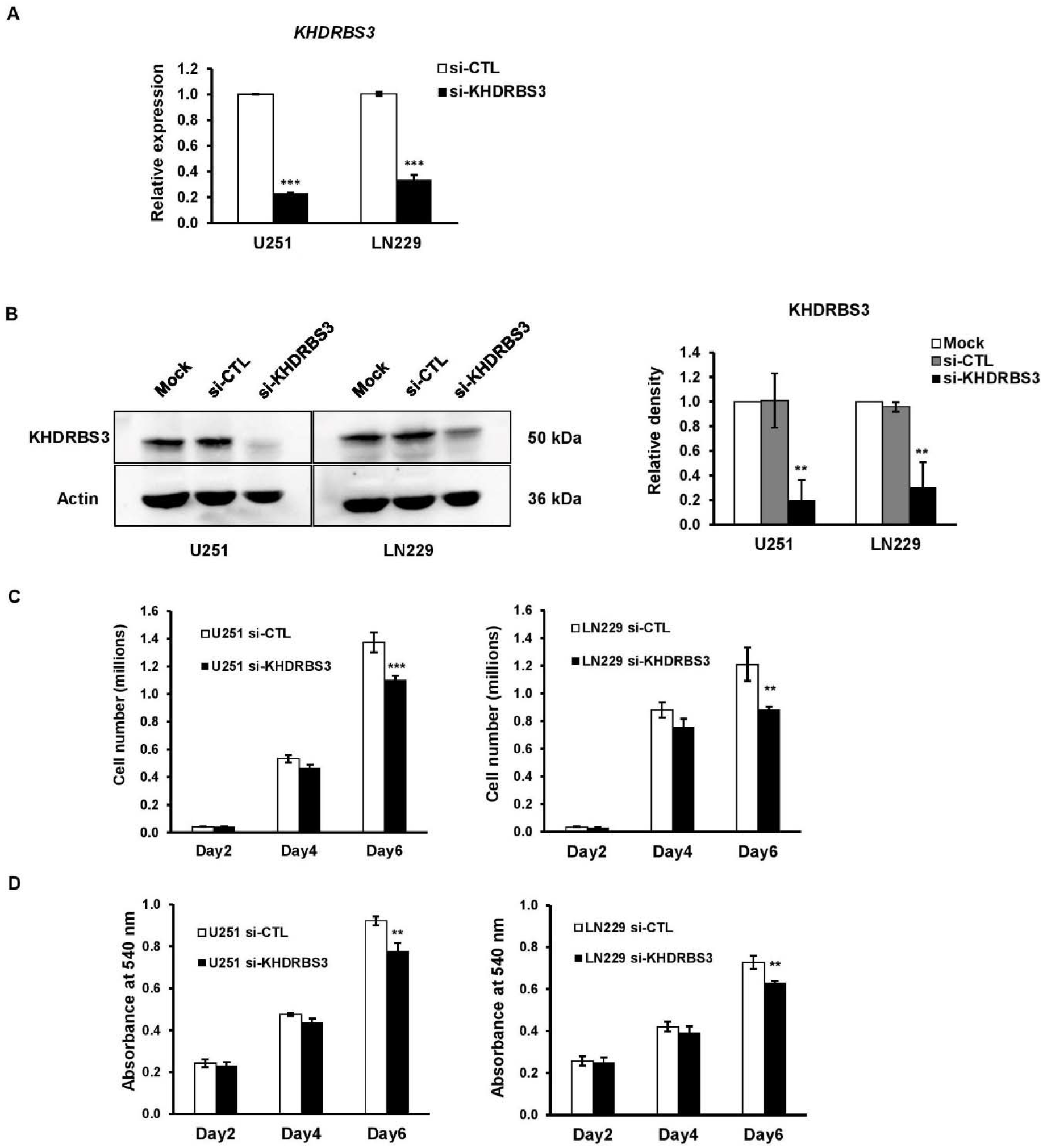
Depletion of KHDRBS3 expression decreased proliferation of U251 and LN229 cells. (A) Realtime PCR analysis of *KHDRBS3* in U251 and LN229 cells which were transfected with 50 nM of negative control siRNA (si-CTL) or *KHDRBS3*-specific siRNA (si-KHDRBS3) for 48 h. *ACTB* was used as internal control. The relative expression value was expressed as the mean ± SEM from three independent experiments. *** p < 0.005. (B) Western blot analysis of KHDRBS3. Band intensities were analyzed and compared using imageJ software. ** p < 0.01 vs mock. (C) si-CTL and si-KHDRBS3 treated U251 and LN229 were grown under monolayer culture up to 6 days. Cell number was counted at the indicated time points. (D) Cell proliferation was determined by MTT assay at the indicated time points. Data is represented as the mean ± SD from three independent experiments. * p < 0.05, ** p < 0.01, ***p<0.005 vs si-CTL.

**Figure 6.**
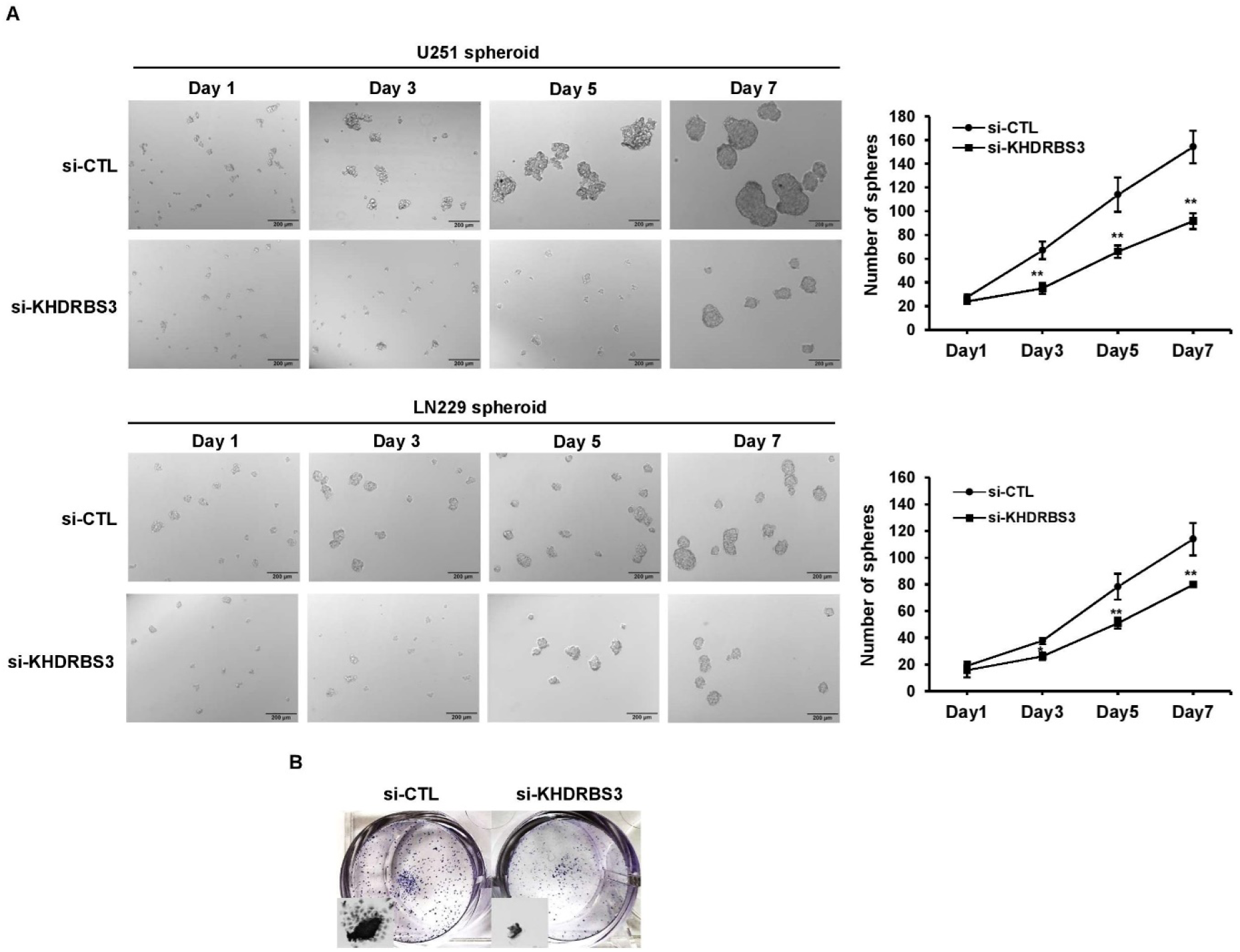
KHDRBS3 facilitates neurosphere formation of U251 and LN229 cells. (A) sphere formation assay was performed in U251 (top) and LN229 (bottom) cells transfected with control (si-CTL) or KHDRBS3 siRNA (si-KHDRBS3). Spheres were photographed by light microscopy at days 3, 5 and 7 (magnification: 10X, scale bar: 200 μm). The number of spheres was counted in and represented as the mean ± SD from three independent experiments. * p < 0.05 and ** p < 0.01 vs si-CTL. (B) A representative image illustrates spheres derived from U251 which were attached to standard culture plates for 6 h in media containing serum, and were stained with crystal violet.

**Figure 7.**
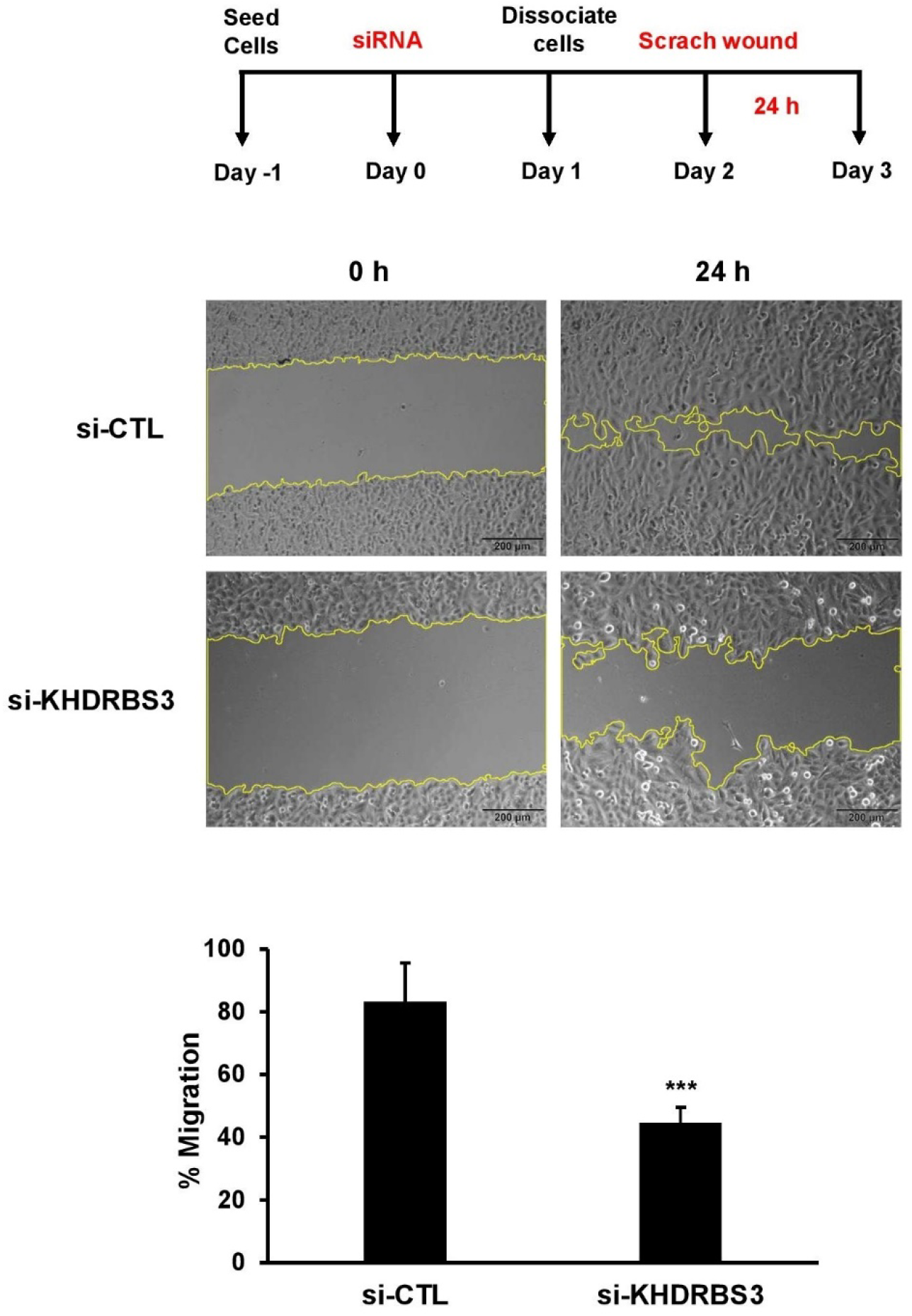
KHDRBS3 promotes cell migration of U251. Wound healing assay of U251 cells transfected with si-CTL or si-KHDRBS3 was performed. Cells were seeded at density of 2.5 x 10^5^ cells/well and incubated in normal culture media for 24 h prior wound scratching. The scratch-wounds were photographed at 0 and 24 h of incubation. Wound areas were measured by Image J software. The histogram shows the percentage of cell migration calculated from the reduction of wound area at 24 h compared with at 0 h. Error bars represent SD from three independent experiments. *** p < 0.005 vs si-CTL.

**Figure 8.**
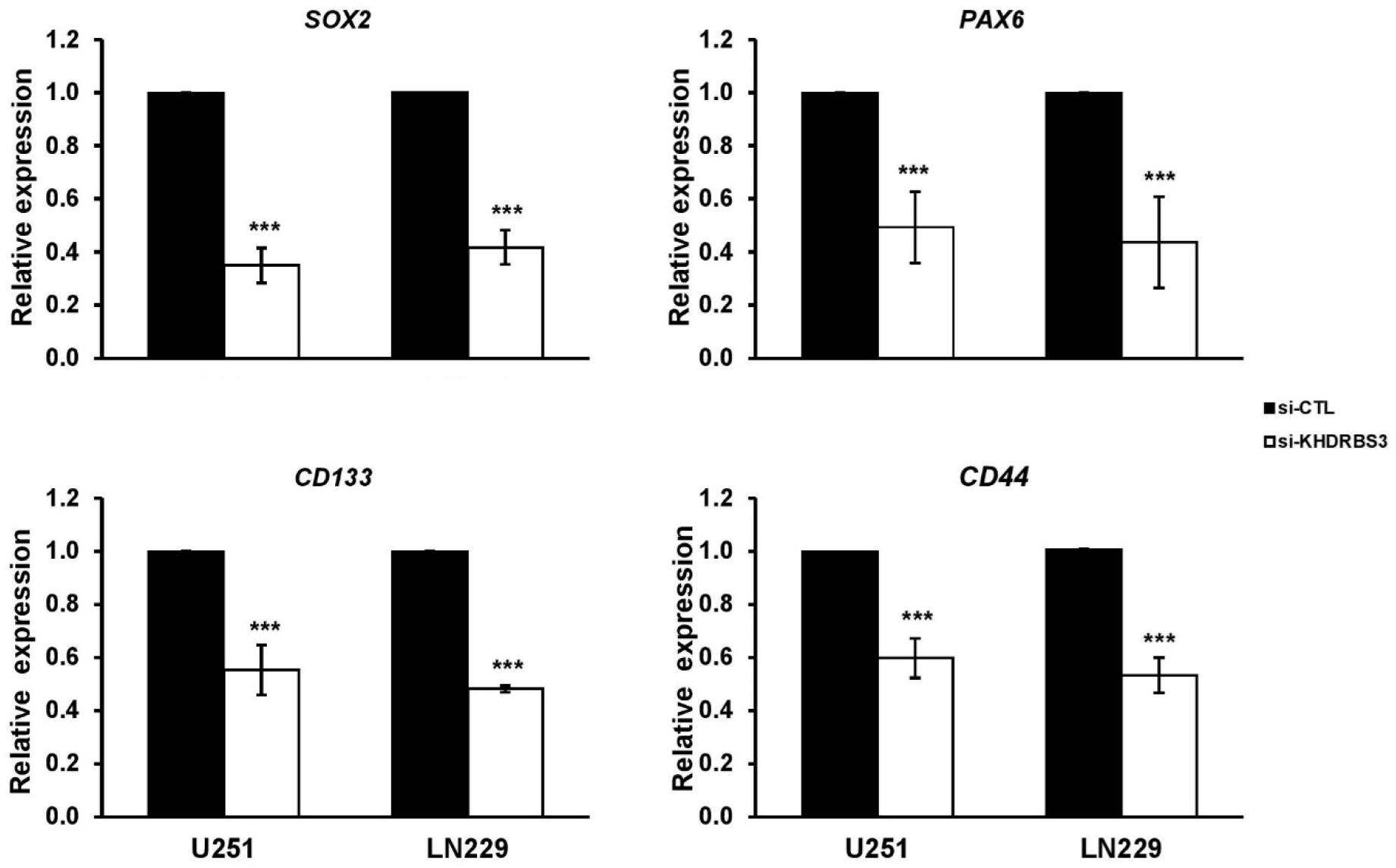
KHDRBS3 facilitates expression of genes involved in cancer stem cells of the glioblastoma cell lines U251 and LN229. Expression of the genes encoding SOX2, PAX6, CD133 and CD44 was determined by realtime PCR in neurospheres derived from U251 and LN229 cells treated with si-CTL or si-KHDRBS3. *ACTB* was used as internal control. The relative value was expressed as the mean ± SD from three independent experiments. ** p < 0.05 and *** p < 0.005 vs si-CTL.

We next sought to test whether KHDRBS3 confers resistance glioblastoma against temozolomide. MTT-based cell viability assay of U251 and LN229 showed that silencing of *KHDRBS3* resulted in a decreased cell viability under temozolomide treatment (Figure 9A). Further, flow cytometry analysis of annexin V/PI staining revealed that KHDRBS3 knockdown cells grown under temozolomide possess an increase in the percentage of annexin V-positive population compared to the control knockdown cells (Figure 9B). Altogether, these data suggest that KHDRBS3 attenuates the chemosensitivity of glioblastoma cells against temozolomide.

**Figure 9.**
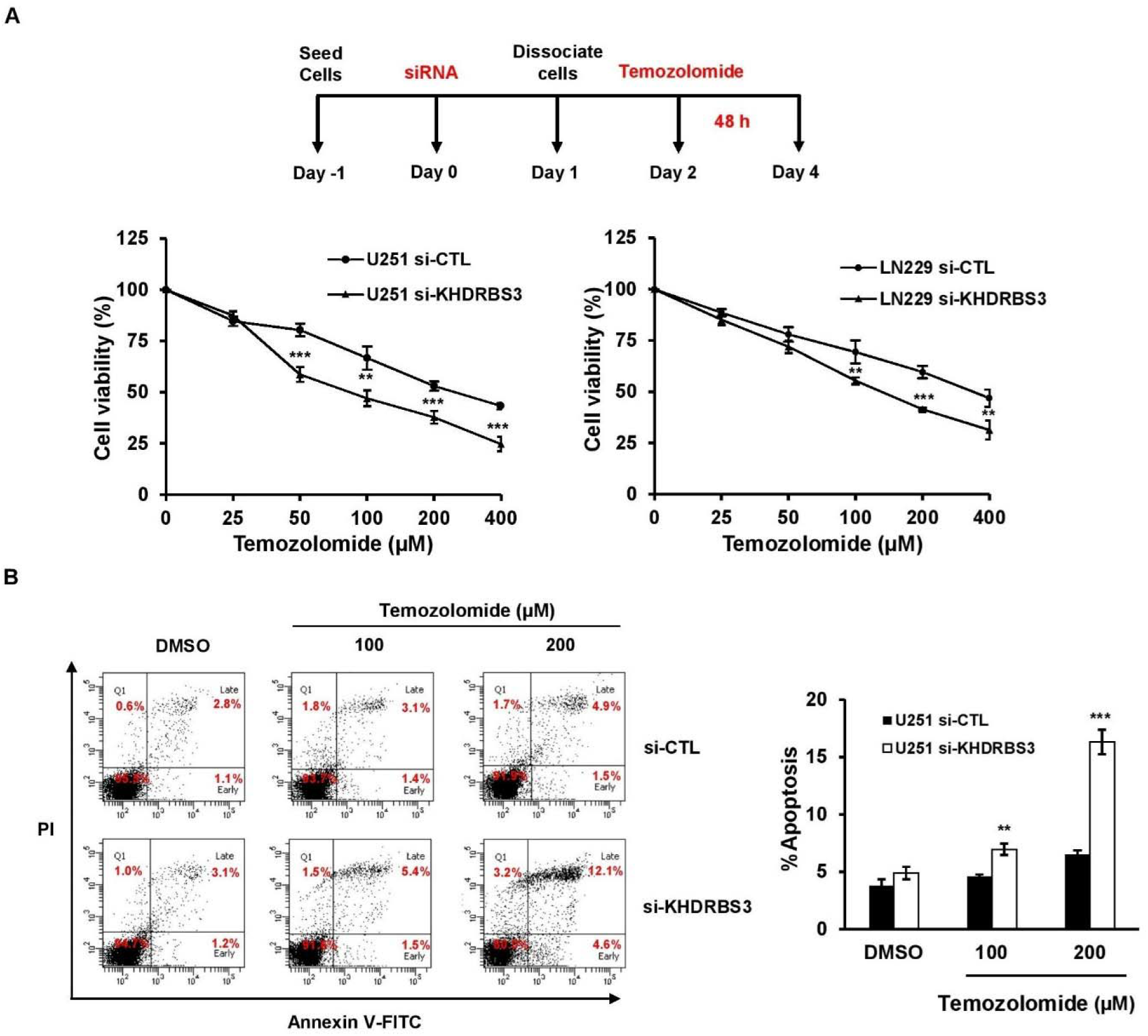
Silencing of KHDRBS3 promotes temozolomide sensitivity in glioblastoma. (A) U251 and LN229 cells transfected with si-CTL or si-KHDRBS3 were cultured in 96-well plates and treated with the indicated concentrations of temozolomide (0-400 µM) for 48 h. The viability of cells treated with temozolomide was determined by MTT assay. (B) Left panel: representative FACS analyses of apoptotic cell death of si-CTL and si-KHDRBS3 treated U251 cells in response to 100 and 200 µM temozolomide. FITC-conjugated Annexin V and PI were used as the early-stage and late-stage apoptotic markers. Right panel: the histogram displayed percentages of apoptotic cells, including early and late apoptotic cells. Error bars represent SD from three independent experiments. ** p < 0.01. *** p < 0.005 compared to si-CTL.

## Discussion

In the present study, we have confirmed that the glioblastoma cell line U251 expresses genes associated with stemness at highest level comparing with other glioblastoma cell lines consistent with previous reports (Hong et al., 2012, Qiang et al., 2009a). We have described the comparative cytoplasmic and nuclear proteomic profiles to identify potential regulators of the cancer stem cell phenotype associated with chemotherapeutic drug resistance in glioblastoma via a neurosphere culture. Pathway analyses revealed that proteins involved in mRNA processing, RNA binding, and spliceosome are significantly enriched in the nuclear proteome of neurosphere cultures of glioblastoma cells when compared with their differentiated counterparts. Our comparative proteomics has confirmed that the anti-apoptotic protein BAG3 is highly expressed in glioblastoma stem-like cells. Interestingly, deletion of BAG3 has been shown to suppress cancer stem cell phenotypes of glioblastoma (Festa et al., 2011, Im et al., 2016). Essentially, we have further identified KHDRBS3 as a protein highly up-regulated in glioblastoma stem-like cells. KHDRBS3 belongs to RNA binding proteins of STAR (signal transduction and the activator of RNA) family containing the KH domain. The STAR family members include KHDRBS1 (also known as Sam68) and KHDRBS2 (also known as SLM1), KHDRBS3 (also known as SLM2), QKI and SF1 which participate in the regulation of selective splicing, export, and stability of RNAs (Vernet and Artzt, 1997). To date, KHDRBS1 and KHDRBS2 have been reported to be associated with human cancers (Benoit et al., 2017, Wang et al., 2015). Yet, the function of KHDRBS3 on stemness and drug resistance in glioblastoma was undescribed.

Up-regulation of KHDRBS3 expression level has been shown for hepatocellular carcinoma (Zhao et al., 2023), medulloblastoma (Lu et al., 2009), ovarian cancer (Wu et al., 2022), and in 5-fluorouracil-resistant gastric cancer cells (Ukai et al., 2020a). Here, we show that the protein KHDRBS3 is up-regulated in glioblastoma stem-like cells including U251, LN229, U87, and M059J spheroids (Figure 4B).

The role of the RNA-binding protein KHDRBS3 in facilitation of stemness has been described in breast (Matsumoto et al., 2018), colon (Ukai et al., 2021), and gastric cancers (Ukai et al., 2020a). In breast cancer stem-like cells, KHDRBS3 controls RNA alternative splicing, and might promote stemness of breast cancer spheroids through an attenuation of anoikis (Matsumoto et al., 2018). Regulation of alternative splicing of a ribosomal gene has been shown to promote cancer stemness in glioblastoma (Larionova et al., 2022). In high-grade glioma, alternative splicing has been identified to activate oncogenic pathways, partly due to the transcription factor REST (Siddaway et al., 2022). In addition, tumorigenesis of glioblastoma has been revealed to associate with several alternative splicing factors such as NONO (Wang et al., 2022b) and DDX3X (Brai et al., 2021). We found that the splicing factor KHDRBS3 facilitates neurosphere formation and expression of genes involved in self-renewal of glioblastoma stem cells (Figures 6 and 8). Both coding and non-coding transcripts have been reported to play a critical role in self-renewal of glioblastoma stem cells. For instance, the histone demethylase KDM1/LSD1 confers self-renewal and temozolomide resistance of glioblastoma stem-like cells (Alejo et al., 2023). Long non-coding RNAs including H19 (Jiang et al., 2016), HIF1A-AS2 (Mineo et al., 2016), SNHG9 (Wang et al., 2022a), and TALNEC2 (Brodie et al., 2017) are also key players in self-renewal of glioblastoma stem-like cells. Since many of these transcripts can be regulated by alternative splicing, as a consequence targeting RNA processing has been proposed as a mean toward novel therapeutics for glioblastoma (Marcelino Meliso et al., 2017).

KHDRBS3 has been shown to facilitate lung metastasis of colorectal and gastric cancers in animal models (Ukai et al., 2020b, Ukai et al., 2021). We show that KHDRBS3 also promotes migration of the glioblastoma cell line U251 (Figure 8). Invasive cancer stem cells can undergo migration and invasion through epithelial-mesenchymal transition (EMT) by which epithelial traits are lost, and mesenchymal phenotypes are intrinsically acquired. Several studies have indicated that the EMT process is associated with stemness and chemoresistance in various cancers such as non-small cell lung, pancreatic, breast, and ovarian cancers (Thiery, 2002). This cell migrating subpopulation is also thought to contribute to chemoresistance and recurrence in glioblastoma (Cho and Kim, 2020, Jung et al., 2016, Lah et al., 2020).

To our knowledge, KHDRBS3 has never been linked to temozolomide resistance. We report here that this is the case for temozolomide resistance in glioblastoma (Figure 9). A number of genes implicated in RNA processing and RNA metabolism play a significant role in resistance against temozolomide in glioblastoma. A genome-wide CRISPR-Cas9 screening has revealed genes implicated in temozolomide resistance in glioblastoma, including FANCA, C19orf40, MCM8, MCM9, and ZC3H7A (MacLeod et al., 2019). Interestingly, the latter is a zinc finger domain-containing RNA-binding protein. Additional RNA-binding proteins found to promote temozolomide resistance include FXR1 (Wei et al., 2022), ADAR3 (Raghava Kurup et al., 2022), NSUN5 (Zhou et al., 2023), RRM2 (Perrault et al., 2023) and MEX3A (Gan et al., 2022). Regarding mRNA alternative splicing, dysregulation of the process is associated with resistance against temozolomide in glioblastoma (Dowdell et al., 2023, Tiek et al., 2022). Among splicing components, the splicing factors FBXO7 and SRSF4 have been identified to facilitate temozolomide resistance of the brain tumor spheroids (Li et al., 2023, Sun et al., 2023). Several transcription and epigenetic factors, which have been recently explored to work closely with their RNA binding partners (Chang and Qi, 2023, Tang et al., 2023), are critical for the drug resistance in glioblastoma. For instance, SOX9 is important not only for self-renewal, but also for temozolomide resistance (Wang et al., 2018). Cell clonogenicity and viability are further reduced in glioblastoma cell lines with silencing of the transcription factor *ATF4* transcript (Lorenz et al., 2021). In addition, the histone lysine demethylase KDM1A/LSD1 confers resistance of temozolomide (Alejo et al., 2023). Apart from coding genes, long non-coding transcripts also play a significant role in aggressiveness of glioblastoma, especially in temozolomide resistance (Li et al., 2022, Roh et al., 2023). Together, our finding has provided an additional evidence, of which KHDRBS3 is factor implicated in resistance of temozolomide. Further studies are crucial to investigate how KHDRBS3 mechanistically regulates the maintenance of self-renewal, malignant properties as well as the drug resistance of glioblastoma. Our results provide a new potential strategy to overcome temozolomide resistance in GBM patients possibly through targeting KHDRBS3-RNA interaction using small molecules (Cai et al., 2017, Disney, 2019, Kovachka et al., 2024).

## Acknowledgements

This research project was supported by CIF Grant, Faculty of Science, Mahidol University. K.S. was supported by Science Achievement Scholarship of Thailand. P.W. was supported by Fundamental Fund, Mahidol University (FF66; FF-067/2566); Program Management Unit for Human Resources and Institutional Development, Research and Innovation (PMU-B; B05F640047); and Mid-Career Scholar Research (National Research Council of Thailand and Mahidol University; N42A650359). We are indebted to WK and PW lab members for their suggestions and comments.

## Author contribution

Conceptualization: KS, PW, WK; Data curation: KS, SK, PW, WK; Formal analysis: KS, PW, WK; Funding acquisition: PW; Methodology: KS, SK, YY, NP; Resources: PW, WK; Manuscript writing, review & editing: KS, SK, PW, WK

---

**Figure S1.**
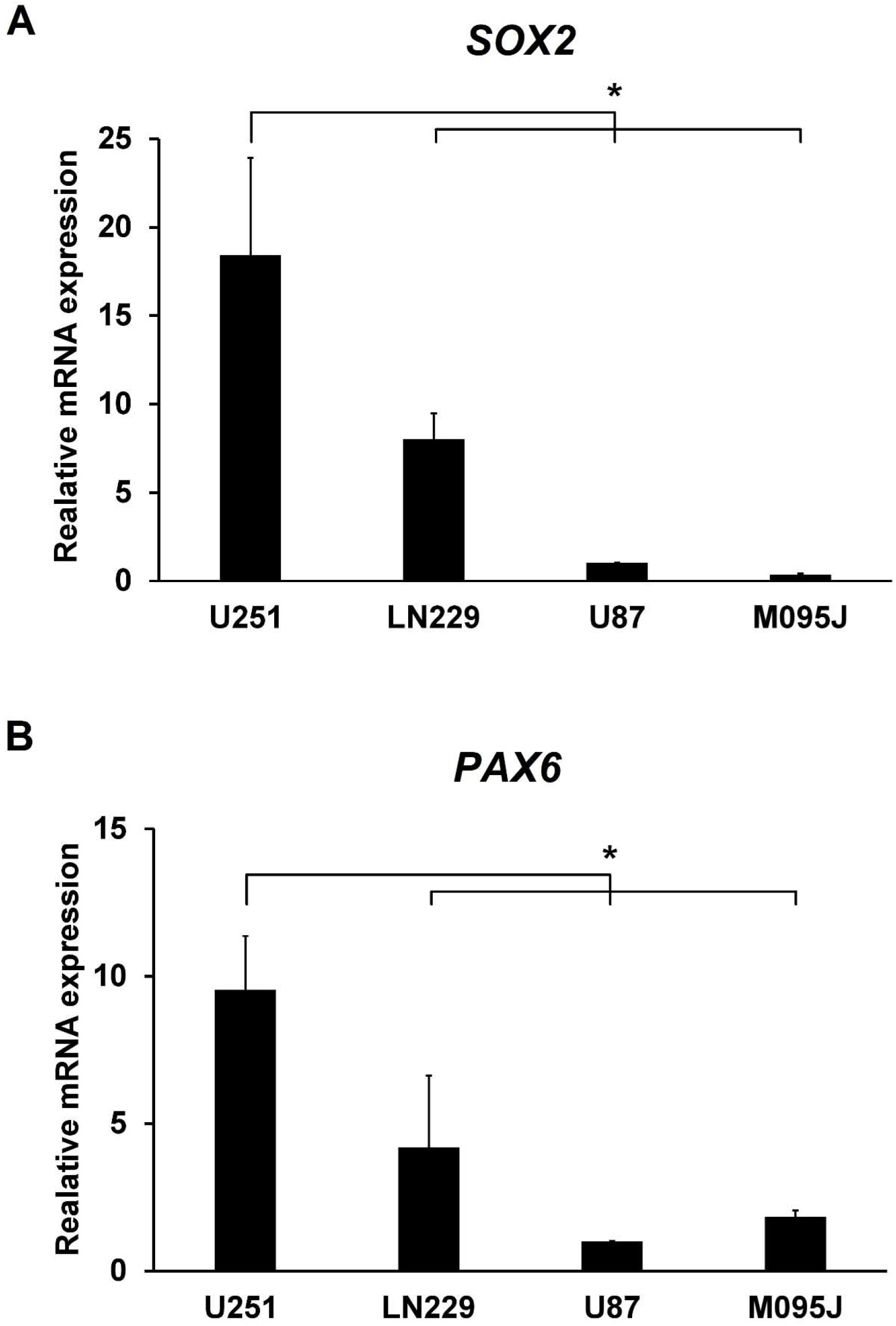
Relative mRNA expression of SOX2 and PAX6 in four glioblastoma cell lines including U251, LN229, U87 and M059J. *ACTB* was used as internal control. The relative value was expressed as the mean ± SD from three independent experiments. * p < 0.01.

**Figure S2.**
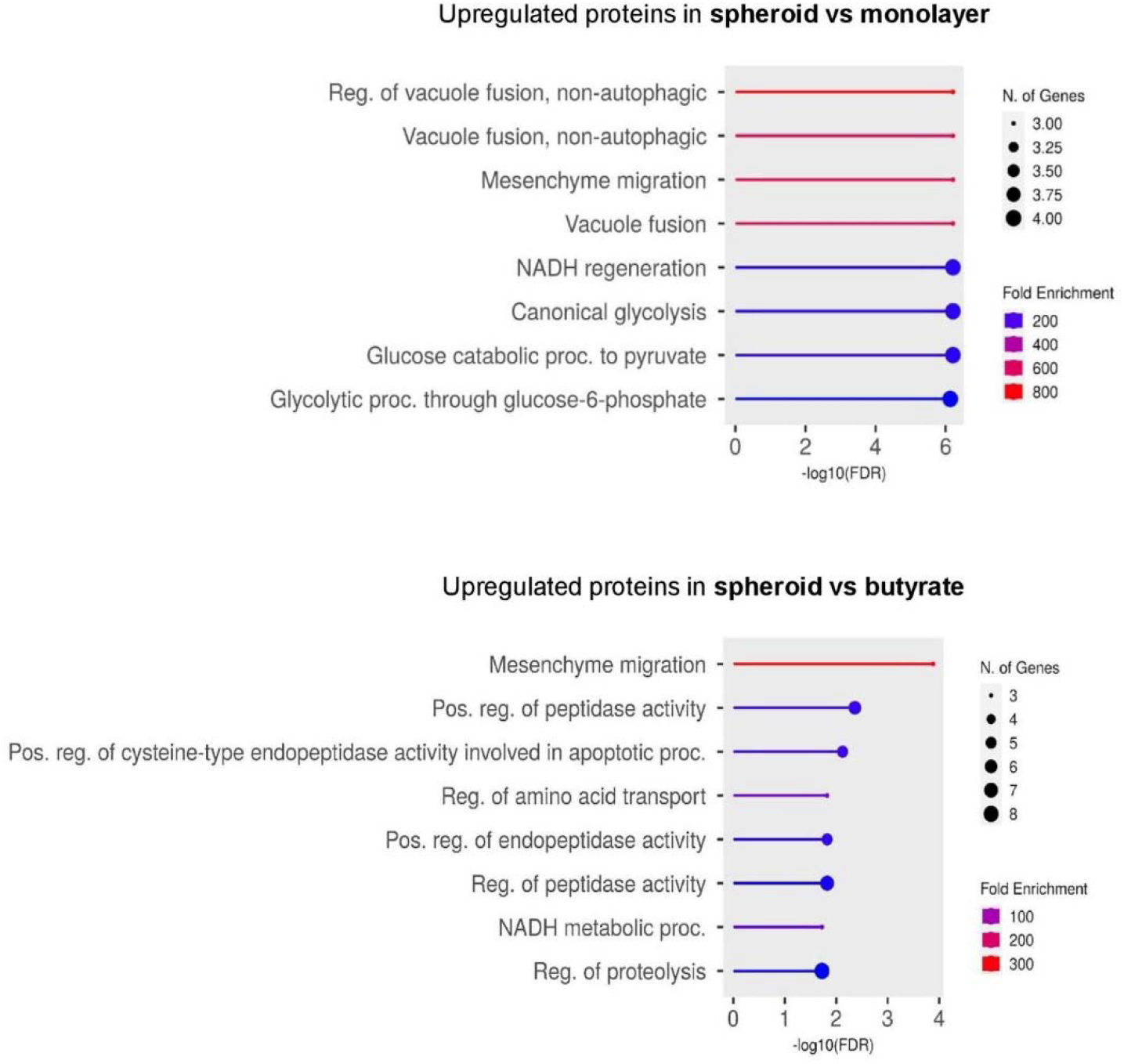
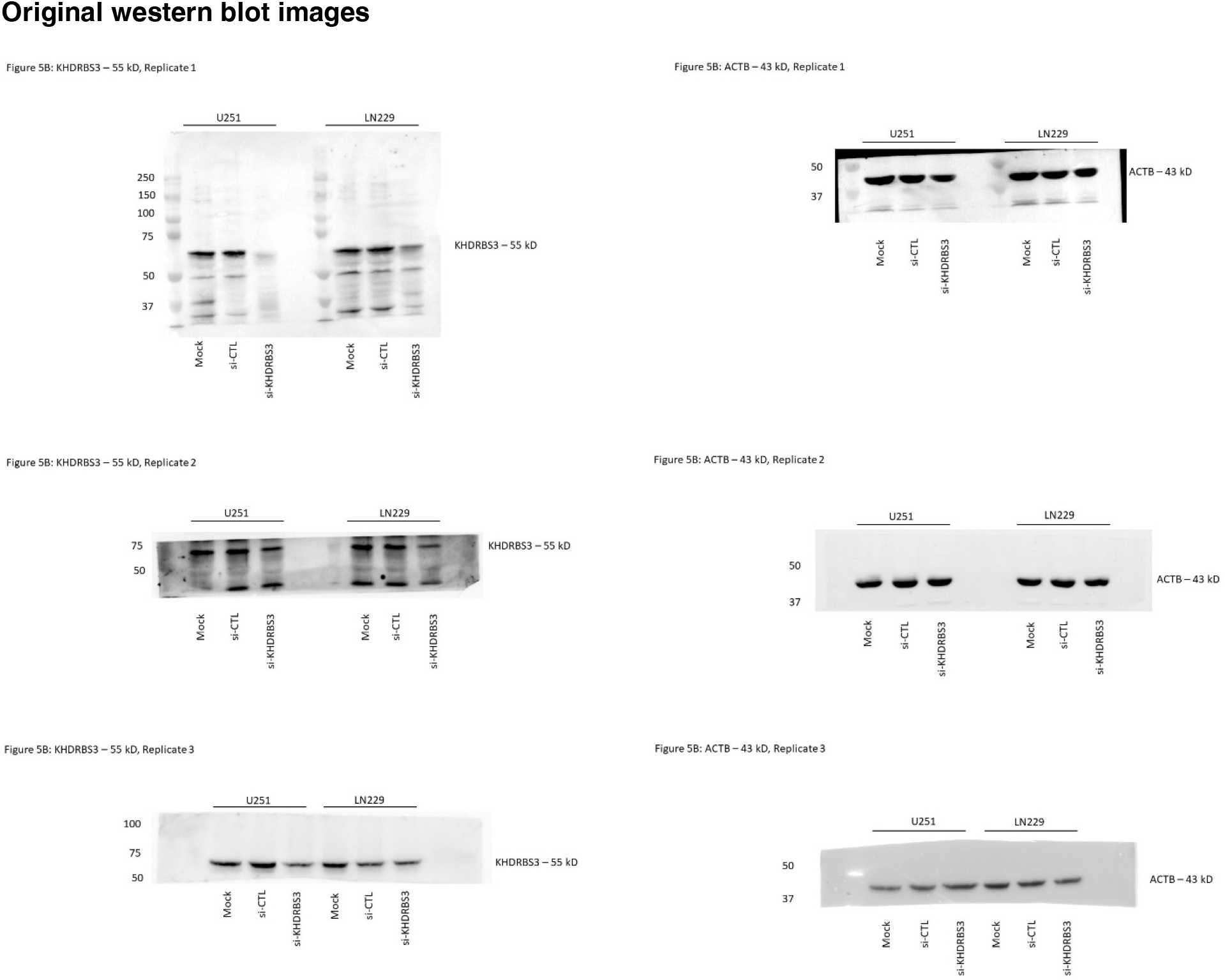
Comparative proteomic analysis between cytoplasmic compartment of spheroid, butyrate and monolayer U251 cells. (top) spheroid vs monolayer, (bottom) spheroid vs butyrate culture cells enriched in the cellular processes. Hierarchical clustering was performed with ShinyGo software v0.76.2. The color chart shows the fold enrichment for each pathway. The size of dots corresponds to the number of genes assigned to each pathway.

## References

1. Alejo S, Palacios BE, Venkata PP, He Y, Li W, Johnson JD, et al. Lysine-specific histone demethylase 1A (KDM1A/LSD1) inhibition attenuates DNA double-strand break repair and augments the efficacy of temozolomide in glioblastoma. Neuro Oncol. 2023;25:1249–61.

2. Auffinger B, Spencer D, Pytel P, Ahmed AU, Lesniak MS. The role of glioma stem cells in chemotherapy resistance and glioblastoma multiforme recurrence. Expert review of neurotherapeutics. 2015;15:741–52.

3. Benoit YD, Mitchell RR, Risueño RM, Orlando L, Tanasijevic B, Boyd AL, et al. Sam68 allows selective targeting of human cancer stem cells. Cell Chemical Biology. 2017;24:833–44. e9.

4. Biserova K, Jakovlevs A, Uljanovs R, Strumfa I. Cancer stem cells: significance in origin, pathogenesis and treatment of glioblastoma. Cells. 2021;10:621.

5. Bradshaw A, Wickremesekera A, Brasch HD, Chibnall AM, Davis PF, Tan ST, et al. Cancer stem cells in glioblastoma multiforme. Frontiers in surgery. 2016;3:48.

6. Brai A, Riva V, Clementi L, Falsitta L, Zamperini C, Sinigiani V, et al. Targeting DDX3X Helicase Activity with BA103 Shows Promising Therapeutic Effects in Preclinical Glioblastoma Models. Cancers (Basel). 2021;13.

7. Brodie S, Lee HK, Jiang W, Cazacu S, Xiang C, Poisson LM, et al. The novel long non-coding RNA TALNEC2, regulates tumor cell growth and the stemness and radiation response of glioma stem cells. Oncotarget. 2017;8:31785–801.

8. Brown DV, Filiz G, Daniel PM, Hollande F, Dworkin S, Amiridis S, et al. Expression of CD133 and CD44 in glioblastoma stem cells correlates with cell proliferation, phenotype stability and intra-tumor heterogeneity. PLoS One. 2017;12:e0172791.

9. Cai W, Xiong Chen Z, Rane G, Satendra Singh S, Choo Z, Wang C, et al. Wanted DEAD/H or Alive: Helicases Winding Up in Cancers. J Natl Cancer Inst. 2017;109.

10. Chang HY, Qi LS. Reversing the Central Dogma: RNA-guided control of DNA in epigenetics and genome editing. Mol Cell. 2023;83:442–51.

11. Chen J, Li Y, Yu T-S, McKay RM, Burns DK, Kernie SG, et al. A restricted cell population propagates glioblastoma growth after chemotherapy. Nature. 2012;488:522–6.

12. Cho Y, Kim YK. Cancer stem cells as a potential target to overcome multidrug resistance. Frontiers in Oncology. 2020;10:764.

13. Disney MD. Targeting RNA with Small Molecules To Capture Opportunities at the Intersection of Chemistry, Biology, and Medicine. J Am Chem Soc. 2019;141:6776–90.

14. Dowdell A, Marsland M, Faulkner S, Gedye C, Lynam J, Griffin CP, et al. Targeting XBP1 mRNA splicing sensitizes glioblastoma to chemotherapy. FASEB Bioadv. 2023;5:211–20.

15. Festa M, Del Valle L, Khalili K, Franco R, Scognamiglio G, Graziano V, et al. BAG3 protein is overexpressed in human glioblastoma and is a potential target for therapy. The American journal of pathology. 2011;178:2504–12.

16. Gan T, Wang Y, Xie M, Wang Q, Zhao S, Wang P, et al. MEX3A Impairs DNA Mismatch Repair Signaling and Mediates Acquired Temozolomide Resistance in Glioblastoma. Cancer Res. 2022;82:4234–46.

17. Hong X, Chedid K, Kalkanis SN. Glioblastoma cell line-derived spheres in serum containing medium versus serum-free medium: A comparison of cancer stem cell properties. International journal of oncology. 2012;41:1693–700.

18. Im C-N, Yun HH, Song B, Youn D-Y, Cui MN, Kim HS, et al. BIS-mediated STAT3 stabilization regulates glioblastoma stem cell-like phenotypes. Oncotarget. 2016;7:35056.

19. Jiang X, Yan Y, Hu M, Chen X, Wang Y, Dai Y, et al. Increased level of H19 long noncoding RNA promotes invasion, angiogenesis, and stemness of glioblastoma cells. J Neurosurg. 2016;2016:129–36.

20. Jiapaer S, Furuta T, Tanaka S, Kitabayashi T, Nakada M. Potential strategies overcoming the temozolomide resistance for glioblastoma. Neurologia medico-chirurgica. 2018;58:405.

21. Jin F, Gao C, Zhao L, Zhang H, Wang H-T, Shao T, et al. Using CD133 positive U251 glioblastoma stem cells to establish nude mice model of transplanted tumor. Brain research. 2011;1368:82–90.

22. Jung EH, Lee HN, Han GY, Kim MJ, Kim CW. Targeting ROR1 inhibits the self-renewal and invasive ability of glioblastoma stem cells. Cell Biochemistry and Function. 2016;34:149–57.

23. Kovachka S, Panosetti M, Grimaldi B, Azoulay S, Di Giorgio A, Duca M. Small molecule approaches to targeting RNA. Nat Rev Chem. 2024.

24. Lah TT, Novak M, Breznik B. Brain malignancies: Glioblastoma and brain metastases. Seminars in cancer biology: Elsevier; 2020. p. 262–73.

25. Larionova TD, Bastola S, Aksinina TE, Anufrieva KS, Wang J, Shender VO, et al. Alternative RNA splicing modulates ribosomal composition and determines the spatial phenotype of glioblastoma cells. Nat Cell Biol. 2022;24:1541–57.

26. Lathia JD, Mack SC, Mulkearns-Hubert EE, Valentim CL, Rich JN. Cancer stem cells in glioblastoma. Genes & development. 2015;29:1203–17.

27. Li S, Chen Y, Xie Y, Zhan H, Zeng Y, Zeng K, et al. FBXO7 Confers Mesenchymal Properties and Chemoresistance in Glioblastoma by Controlling Rbfox2-Mediated Alternative Splicing. Adv Sci (Weinh). 2023;10:e2303561.

28. Li S, Xie X, Peng F, Du J, Peng C. Regulation of temozolomide resistance via lncRNAs: Clinical and biological properties of lncRNAs in gliomas (Review). Int J Oncol. 2022;61.

29. Lorenz NI, Sittig ACM, Urban H, Luger AL, Engel AL, Münch C, et al. Activating transcription factor 4 mediates adaptation of human glioblastoma cells to hypoxia and temozolomide. Sci Rep. 2021;11:14161.

30. Lu Y, Ryan SL, Elliott DJ, Bignell GR, Futreal PA, Ellison DW, et al. Amplification and overexpression of Hsa-miR-30b, Hsa-miR-30d and KHDRBS3 at 8q24. 22-q24. 23 in medulloblastoma. PloS one. 2009;4:e6159.

31. MacLeod G, Bozek DA, Rajakulendran N, Monteiro V, Ahmadi M, Steinhart Z, et al. Genome-Wide CRISPR-Cas9 Screens Expose Genetic Vulnerabilities and Mechanisms of Temozolomide Sensitivity in Glioblastoma Stem Cells. Cell Rep. 2019;27:971–86.e9.

32. Marcelino Meliso F, Hubert CG, Favoretto Galante PA, Penalva LO. RNA processing as an alternative route to attack glioblastoma. Hum Genet. 2017;136:1129–41.

33. Matsumoto Y, Itou J, Sato F, Toi M. SALL4-KHDRBS3 network enhances stemness by modulating CD 44 splicing in basal-like breast cancer. Cancer medicine. 2018;7:454–62.

34. McNamara MG, Lwin Z, Jiang H, Chung C, Millar B-A, Sahgal A, et al. Conditional probability of survival and post-progression survival in patients with glioblastoma in the temozolomide treatment era. Journal of neuro-oncology. 2014;117:153–60.

35. Mineo M, Ricklefs F, Rooj AK, Lyons SM, Ivanov P, Ansari KI, et al. The Long Non-coding RNA HIF1A-AS2 Facilitates the Maintenance of Mesenchymal Glioblastoma Stem-like Cells in Hypoxic Niches. Cell Rep. 2016;15:2500–9.

36. Norden AD, Reardon DA, Wen PC. Primary central nervous system tumors: Pathogenesis and therapy: Springer Science & Business Media; 2010.

37. Pang R, Law WL, Chu AC, Poon JT, Lam CS, Chow AK, et al. A subpopulation of CD26+ cancer stem cells with metastatic capacity in human colorectal cancer. Cell stem cell. 2010;6:603–15.

38. Perrault EN, Shireman JM, Ali ES, Lin P, Preddy I, Park C, et al. Ribonucleotide reductase regulatory subunit M2 drives glioblastoma TMZ resistance through modulation of dNTP production. Sci Adv. 2023;9:eade7236.

39. Phi LTH, Sari IN, Yang YG, Lee SH, Jun N, Kim KS, et al. Cancer Stem Cells (CSCs) in Drug Resistance and their Therapeutic Implications in Cancer Treatment. Stem Cells Int. 2018;2018:5416923.

40. Putavet DA, de Keizer PL. Residual disease in glioma recurrence: A dangerous liaison with senescence. Cancers. 2021;13:1560.

41. Qiang L, Yang Y, Ma Y-J, Chen F-H, Zhang L-B, Liu W, et al. Isolation and characterization of cancer stem like cells in human glioblastoma cell lines. Cancer letters. 2009a;279:13–21.

42. Qiang L, Yang Y, Ma YJ, Chen FH, Zhang LB, Liu W, et al. Isolation and characterization of cancer stem like cells in human glioblastoma cell lines. Cancer Lett. 2009b;279:13–21.

43. Raghava Kurup R, Oakes EK, Vadlamani P, Nwosu O, Danthi P, Hundley HA. ADAR3 activates NF-κB signaling and promotes glioblastoma cell resistance to temozolomide. Sci Rep. 2022;12:13362.

44. Roh J, Im M, Kang J, Youn B, Kim W. Long non-coding RNA in glioma: novel genetic players in temozolomide resistance. Anim Cells Syst (Seoul). 2023;27:19–28.

45. Siddaway R, Milos S, Vadivel AKA, Dobson THW, Swaminathan J, Ryall S, et al. Splicing is an alternate oncogenic pathway activation mechanism in glioma. Nat Commun. 2022;13:588.

46. Sun Y, Liu X, Wu Z, Wang X, Zhang Y, Yan W, et al. SRSF4 Confers Temozolomide Resistance of Glioma via Accelerating Double Strand Break Repair. J Mol Neurosci. 2023;73:259–68.

47. Tang J, Wang X, Xiao D, Liu S, Tao Y. The chromatin-associated RNAs in gene regulation and cancer. Mol Cancer. 2023;22:27.

48. Thiery JP. Epithelial–mesenchymal transitions in tumour progression. Nature reviews cancer. 2002;2:442–54.

49. Tiek DM, Erdogdu B, Razaghi R, Jin L, Sadowski N, Alamillo-Ferrer C, et al. Temozolomide-induced guanine mutations create exploitable vulnerabilities of guanine-rich DNA and RNA regions in drug-resistant gliomas. Sci Adv. 2022;8:eabn3471.

50. Tomar MS, Kumar A, Srivastava C, Shrivastava A. Elucidating the mechanisms of Temozolomide resistance in gliomas and the strategies to overcome the resistance. Biochimica et Biophysica Acta (BBA)-Reviews on Cancer. 2021;1876:188616.

51. Ukai S, Honma R, Sakamoto N, Yamamoto Y, Pham QT, Harada K, et al. Molecular biological analysis of 5-FU-resistant gastric cancer organoids; KHDRBS3 contributes to the attainment of features of cancer stem cell. Oncogene. 2020a;39:7265–78.

52. Ukai S, Honma R, Sakamoto N, Yamamoto Y, Pham QT, Harada K, et al. Molecular biological analysis of 5-FU-resistant gastric cancer organoids; KHDRBS3 contributes to the attainment of features of cancer stem cell. Oncogene. 2020b;39:7265–78.

53. Ukai S, Sakamoto N, Taniyama D, Harada K, Honma R, Maruyama R, et al. KHDRBS3 promotes multi-drug resistance and anchorage-independent growth in colorectal cancer. Cancer Sci. 2021;112:1196–208.

54. Vernet C, Artzt K. STAR, a gene family involved in signal transduction and activation of RNA. Trends in Genetics. 1997;13:479–84.

55. Wang CJ, Chao CR, Zhao WF, Liu HM, Feng JS, Cui YX. Long noncoding RNA SNHG9 facilitates growth of glioma stem-like cells via miR-326/SOX9 axis. J Gene Med. 2022a;24:e3334.

56. Wang L, Tian H, Yuan J, Wu H, Wu J, Zhu X. CONSORT: Sam68 is directly regulated by miR-204 and promotes the self-renewal potential of breast cancer cells by activating the Wnt/beta-catenin signaling pathway. Medicine. 2015;94.

57. Wang X, Han M, Wang S, Sun Y, Zhao W, Xue Z, et al. Targeting the splicing factor NONO inhibits GBM progression through GPX1 intron retention. Theranostics. 2022b;12:5451–69.

58. Wang Z, Xu X, Liu N, Cheng Y, Jin W, Zhang P, et al. SOX9-PDK1 axis is essential for glioma stem cell self-renewal and temozolomide resistance. Oncotarget. 2018;9:192–204.

59. Wei Y, Duan S, Gong F, Li Q. The RNA-binding protein fragile-X mental retardation autosomal 1 (FXR1) modulates glioma cells sensitivity to temozolomide by regulating ferroptosis. Biochem Biophys Res Commun. 2022;603:153–9.

60. Wongtrakoongate P. Epigenetic therapy of cancer stem and progenitor cells by targeting DNA methylation machineries. World J Stem Cells. 2015;7:137–48.

61. Wu W, Klockow JL, Zhang M, Lafortune F, Chang E, Jin L, et al. Glioblastoma multiforme (GBM): An overview of current therapies and mechanisms of resistance. Pharmacological Research. 2021;171:105780.

62. Wu X, Qiu L, Feng H, Zhang H, Yu H, Du Y, et al. KHDRBS3 promotes paclitaxel resistance and induces glycolysis through modulated MIR17HG/CLDN6 signaling in epithelial ovarian cancer. Life Sci. 2022;293:120328.

63. Zhao M, Zhang Y, Li L, Liu X, Zhou W, Wang C, et al. KHDRBS3 accelerates glycolysis and promotes malignancy of hepatocellular carcinoma via upregulating 14-3-3ζ. Cancer Cell Int. 2023;23:244.

64. Zhou J, Kong YS, Vincent KM, Dieters-Castator D, Bukhari AB, Glubrecht D, et al. RNA cytosine methyltransferase NSUN5 promotes protein synthesis and tumorigenic phenotypes in glioblastoma. Mol Oncol. 2023;17:1763–83.

